# A novel retinoic acid analog, 4-Amino-2-Trifluoromethyl-Phenyl Retinate (ATPR) triggers differentiation and is effective in the treatment of Acute Promyelocytic Leukemia

**DOI:** 10.1101/2020.07.09.194928

**Authors:** Jing Bao, Yan Du, Lan-lan Li, Liang Xia, Fei-hu Chen

**Affiliations:** School of Pharmacy, Anhui Province Key Laboratory of Major Autoimmune Diseases, Anhui Institute of Innovative Drugs, Anhui Medical University, Hefei 230032, China; Institute for Liver Disease of Anhui Medical University, Anhui Medical University, Hefei 230032, China; Department of Hematology, the First Affiliated Hospital of Anhui Medical University, Hefei, Anhui, China

**Keywords:** Acute promyelocytic leukemia (APL), All-trans retinoic acid (ATRA), 4-Amino-2-Trifluoromethyl-Phenyl Retinate (ATPR), PI3K/AKT signaling

## Abstract

Acute promyelocytic leukemia (APL), form RARα fusion genes and proteins is one of the most prevalent forms of leukemia. 4-Amino-2-Trifluoromethyl-Phenyl Retinate (ATPR), a derivative of all-trans-retinoic acid (ATRA), is of potent functions in the induce of cell differentiation, growth arrest, and apoptosis. Nowadays, we aimed to investigate the therapeutic effect of ATPR on this APL pathological model, and whether the mechanism involves PI3K/AKT, ERK and Notch signaling. We established a human xenograft mouse model using NB4 cells and found that ATPR significantly increased the protein concentration in the CD11b and suppressed the PI3K/AKT signaling and activated the ERK and Notch signaling in tumor tissue. Collectively, these data suggest that ATPR shows antileukemic effects by regulating PI3K/AKT, ERK and Notch signaling.

## 1. Introduction

Acute promyelocytic leukemia (APL), once a severe fulminant disease with a high-risk of bleeding tendency. It is driven by a 15;17 chromosome translocation and the formation of leukemogenic fusion protein promyelocytic leukemia–retinoic acid a receptor (PML-RARα)[1]. As an abnormal transcript, PML/RARα has a pivotal role in the pathogenesis of APL, it impairs formation of functional PML nuclear bodies and triggers the tumor phenotype by recruiting the co-repressor histone deacetylase (HDAC)-complex and methyltransferases[2]. The therapy of combining all-trans-retinoic acid (ATRA) and arsenic trioxide (ATO) is the key strategy of the APL treatment [3]. However, there are still about 20–30% patients relapsing, and they are more likely to develop the cardiactoxicity, drug resistance and retinoic acid syndrome (RAS), have been identified as a clinically significant problem[4, 5]. This highlights the pressing need for the other highly effective and safe drugs to help develop the differentiation therapy for APL.

The College of Pharmacy, Anhui Medical University (Hefei, China) designed and synthesized 4-Amino-2-Trifluoromethyl-Phenyl Retinate (ATPR), a derivative of ATRA. Our previous studies revealed that ATPR could exert the superior antitumor activity than ATRA on human gastric cancer[6], hepatocellular carcinoma[7], gastric carcinoma[8], breast cancer[9], especially acute promyelocytic leukemia[10]. There have been studies that indicated the stimulating effect of ATPR on the differentiation *in vitro*, it prompts the therapeutic effects in APL patients. Animal experiment is significant to determine the therapeutic effect of ATPR *in vivo* beforehand, although these animal models would be limited and would not reflect the real situation in patients completely.

It is not like solid tumor to establish tumor-bearing animal model by the xenotransplantation, hematological malignancies with dispersivity and generalized characteristics are difficult to propagate [11–13]. In recent years, a lot of human malignant hematopoietic cells, such as acute lymphoblastic leukemia (ALL), chronic myeloid leukemia (CML), and multiple myeloma (MM), have been successfully engrafted in nude mice[5, 14]. Severe immune-deficiency strain NCG is established by CRISPR/Cas9 technology. Prkdc and Il2rg genes are knocked out on NOD/ShiLtJNju background. Lack of T, B, NK cells. It is a ideal strain for cancer modeling and studies in immune-oncology and efficacy testing[15, 16].

Here, we showed the human xenograft mouse model established by using NB4 cells and confirmed the therapeutic effect of ATPR on this APL pathological model. Furthermore, ATPR may have latent capacity to be applied in clinical as a new chemotherapy.

## 2. Materials and methods

### 2.1 Materials

ATPR was obtained from the college of Pharmacy from Anhui Medical University (Anhui, China), with 99.66% purification (Figure 1A). ATRA was purchased from Sigma (USA).

**Figure 1.**
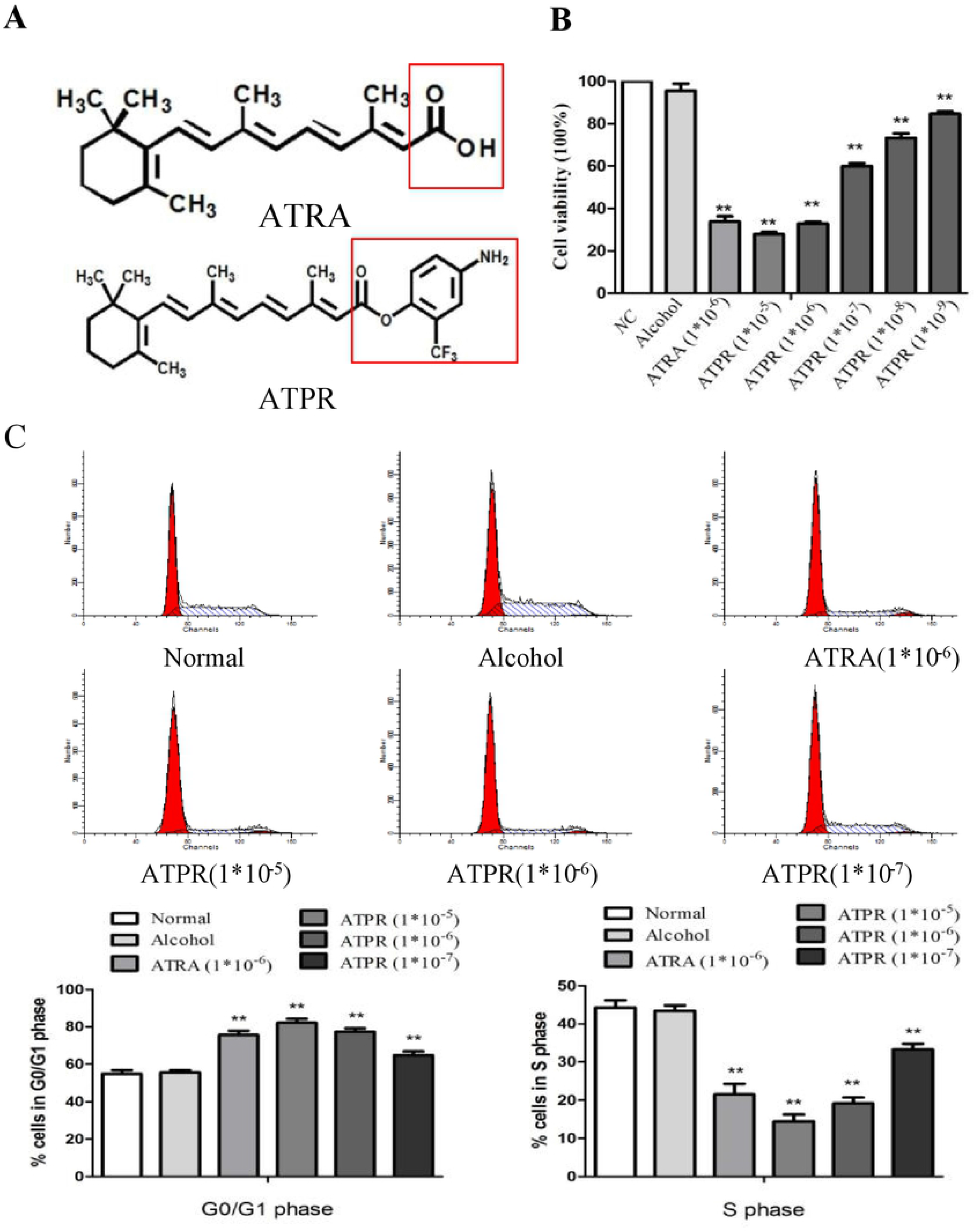
Effects of ATPR on the cell viability and cell cycle in NB4 cells. A. Chemical structure of ATPR, a novel all-trans retinoic acid (ATRA) derivative, designed and synthesized by our team. B. Concentration-dependent inhibition of cellular viability of ATRA (1*10^-6^ mol/L), ATPR (1*10^-9^, 1*10^-8^, 1*10^-7^, 1*10^-6^, 1*10^-5^ mol/L) on NB4 cells as measured by CCK-8 assay. \**p*<0.05, \*\**p*<0.01 versus control group. C. NB4 cells were treated with ATPR (1*10^-7^, 1*10^-6^, 1*10^-5^ mol/L) and ATRA (1*10^-6^ mol/L). The cell cycle distribution was analyzed by flow cytometry. \**p*<0.05, \*\**p*<0.01 versus control group.

### 2.2 Cells and cell culture

The APL cell line NB4 was purchased from Genechem Co.Ltd. (Shanghai, China). NB4 cells were maintained in suspension in RPMI-1640 medium (Hyclone) containing 8% FBS (Biological Industries), 100 U/ml penicillin and 100 g/ml streptomycin in 5% CO2, 37 °C incubator.

### 2.3 Transplantation of NB4 cells into NCG mice

NCG mice were purchased from the model animal research institute (Nanjing, China), and the animal experiment was approved by the ethics committee as well as the Animal Care and Use Committee in Anhui Medical University (Hefei, China). The mice had been quarantined for a week before we conducted the experiments in SPF laboratory animal room. The APL mice models were built by inoculating with 100 μl of NB4 cells (5×10^6^) diluted with the matrigel and PBS mixture (1:1) on the right shoulder of each mouse. The tumor growth was observed and measured daily, when the tumor volume exceeded 100mm^3^, mice models were randomly divided into two groups and accepted different treatments: experimental group received ATPR (10 mg/kg) per day while normal control group dosed equal normal solvent. Continuous administration for one week, all mice were observed and recorded changes in their weight and tumor volume every day. (volume=(length×width2)/2). After a week, the mice were killed to prepare the tumors tissue sections for H&E staining and immunohistochemical analysis. All experiments were repeated at least for three times.

All experiments were designed in strict conformity with the NIH guidelines and were approved by the institutional ethics guidelines for laboratory animal care and approved by the Animal Care and Use Committee in Anhui Medical University.

### 2.4 Hematological tests

Peripheral venous blood samples were collected in each pre-determined time and hematological parameters such as erythrocyte, leukocyte, and hemoglobin (Hb) were counting. 20ul blood was inhaled through a micropipette from the tail vein of each mice and 180ul sulfolyser (Sysmex) were added. All sample testing was determined in the Second Affiliated Hospital of Anhui Medical University using the same automated blood cell analyzer (XT-4000i, Sysmex, Japan).

### 2.5 Western blotting

Tumor tissues were lysed in RIPA lysis buffer for 30 min and then centrifuged at 12,000rpm for 40 min to collect the supernatant. BCA protein assay kit (Boster, China) was used to measure the protein concentrations. The protein samples (20 or 40μg) were separated by 8% and 12% sodium dodecyl sulfate-polyacrylamide gel electrophoresis (SDS-PAGE) before transferred to PVDF membranes (Millipore, Billerica, MA, USA). The membranes were blocked with 5% skim milk compound with TBST for 3h and incubated with corresponding primary antibodies and appropriate secondary antibodies before the protein bands were detected with ECL-chemiluminescent kit (ECLplus, Thermo Scientific). Antibodies against human CD11b, p-AKT, AKT, PI3K, p-ERK 1/2, ERK 1/2, Notch 1 and Hes 1 (Abcam) were diluted at 1:1000 dilution and anti-β-actin (ZSGB-BIO, China) were used at a concentration of 1:300. Image J software was applied to analyze the immunoblotting images in quantitative density. The experiment was repeated for three times.

### 2.6 Statistical analysis

Experimental data were expressed as means ±SD of three determinations. An analysis of variance (ANOVA), Student’s t-test and Bonferroni’s test were performed for statistical analysis. *P*<0.05 was considered statistically significant.

## 3. Result

### 3.1 Effects of ATPR on the cell viability and cell cycle in NB4 cells

MTT assay was used to evaluate the distinct cellular viabilities of NB4 cells influenced by ATPR with different concentrations (1*10^-9^, 1*10^-8^, 1*10^-7^, 1*10^-6^, 1*10^-5^ mol/L), contrasting with ATRA (1*10^-6^ mol/L) (Figure 1B). As shown in figure, growth-inhibitory effects of ATPR on NB4 cells were dose-dependent and the IC50 of ATPR is 3.7*10^-6^ mol/L. According to the pre-experiment, the concentration and treating time of ATPR were decided (1*10^-6^ mol/L, 72h). FCM (flow cytometry) was used to explore how ATPR effect the proliferation of NB4 cells. As shown in Fig.1C, a dose-dependent accumulation of cells in G0/G1 phase but the reductions of cells percentage in S phase were observed after ATPR treatment. It indicated that NB4 cells were arrested in G0/G1 phase by ATPR.

### 3.2 Effects of ATPR on the cell differentiation in NB4 cells

CD11b was the myeloid differentiation markers. The cells were examined by FCM with antibodies against CD11b. Interestingly, FCM suggested that ATPR markedly upregulated the expression level of CD11b in NB4 cells in comparison with control group (Figure 2). These data indicated that ATPR might have a big impact on inducing NB4 cells differentiation and growth arrest.

**Figure 2.**
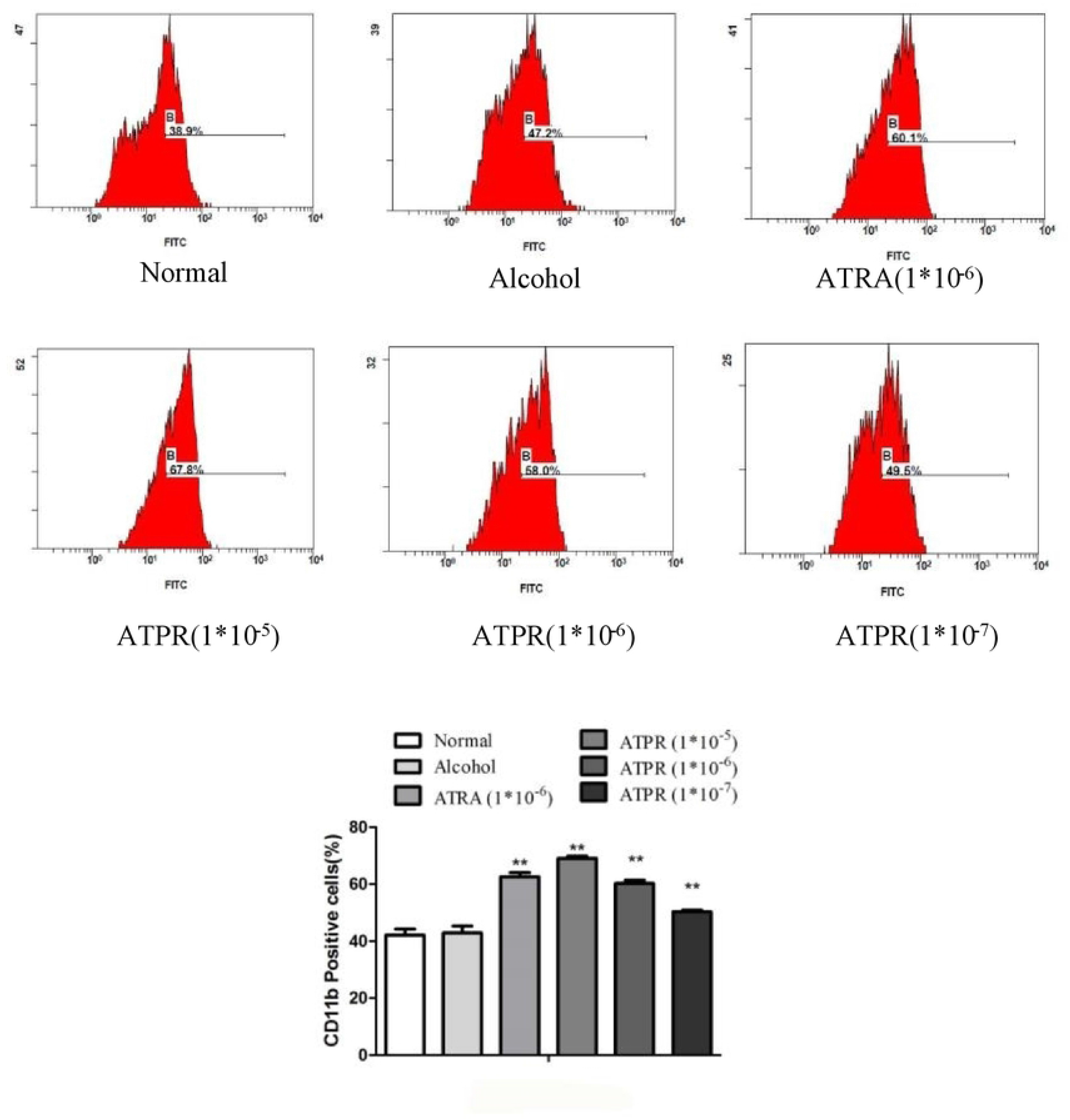
Effects of ATPR on the cell differentiation in NB4 cells. NB4 cells were treated with ATPR (1*10^-7^, 1*10^-6^, 1*10^-5^ mol/L) and ATRA (1*10^-6^ mol/L). CD11b expression was analyzed by flow cytometry. \**p*<0.05, \*\**p*<0.01 versus control group.

### 3.3 ATPR shows antileukemic effects through the signaling pathway *in vitro*

To explain the mechanism of ATPR on the NB4 cells, the PI3K/AKT, ERK and Notch signaling pathways may be considered to participate because these have been shown to regulate the APL-cells differentiation, proliferation, and apoptosis *in vitro*. We found that ATPR reduced the expressions of p-Akt, PI3K and enhanced the levels of p-ERK1/2, Notch1 and Hes1 (Fig. 3). These results furtherly confirmed that ATPR shows antileukemic effects through the PI3K/AKT, ERK, and Notch signaling pathways.

**Figure 3.**
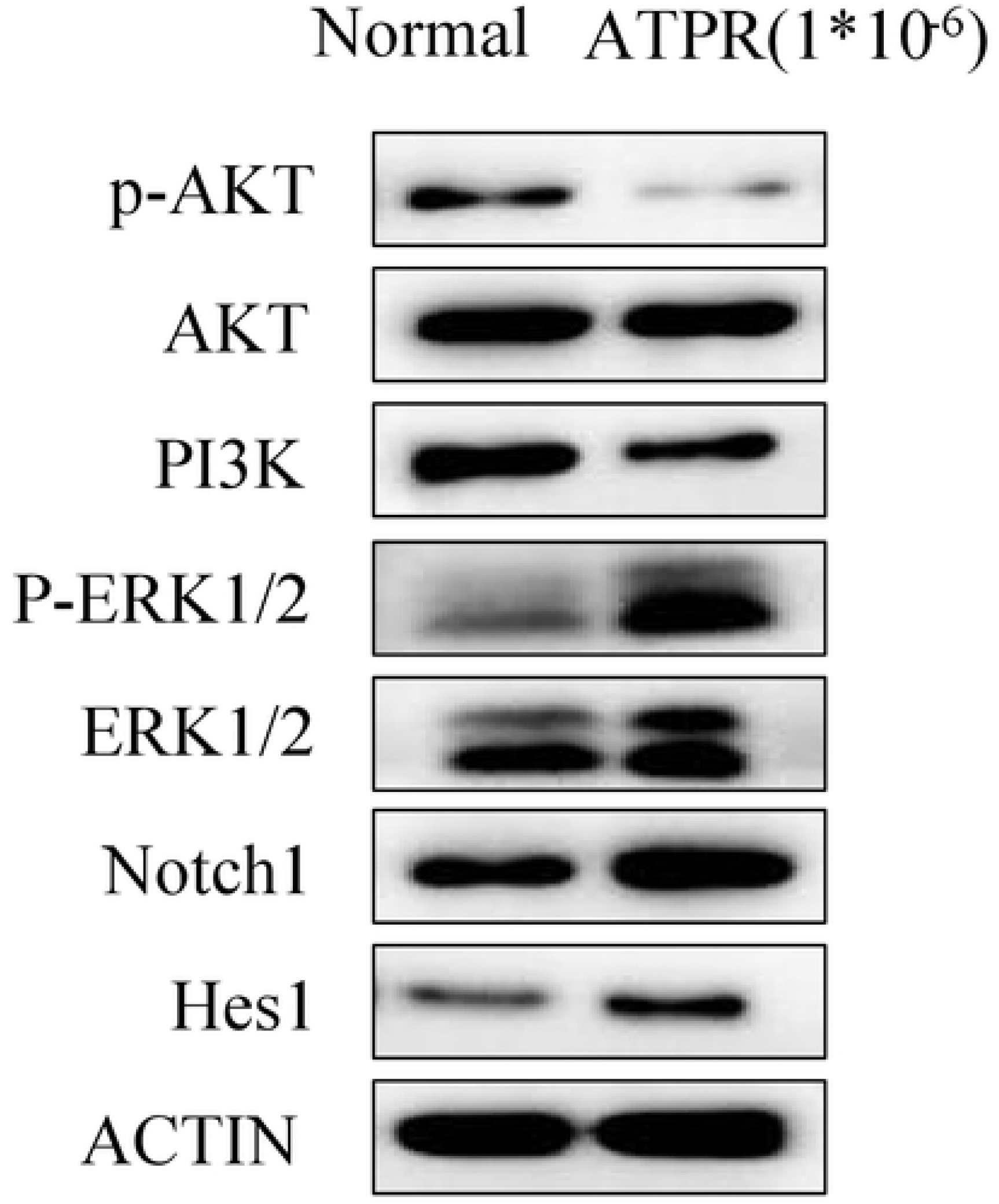
ATPR shows antileukemic effects *in vitro*. Western blotting was performed to detect the expression levels of p-AKT, AKT, PI3K, p-ERK1/2, ERK1/2, Notch1 and Hes1. \**p*<0.05, \*\**p*<0.01 versus control group.

### 3.4 ATPR shows antileukemic effects through the signaling pathway *in vivo*

To explain the mechanism of ATPR on the mice, the PI3K/AKT, ERK and Notch signaling pathways may be considered to participate. Compared with vehicle group, ATPR group showed the growth suppression on subcutaneous tumor in NCG mice (Fig. 4A, B). Furthermore, we found that ATPR-treated mice reduced the expressions of p-Akt, PI3K and enhanced the levels of p-ERK1/2, Notch1 and Hes1 (Fig. 4C) in comparison to vehicle-treated mice. These results furtherly confirmed that ATPR shows antileukemic effects through the PI3K/AKT, ERK, and Notch signaling pathways.

**Figure 4.**
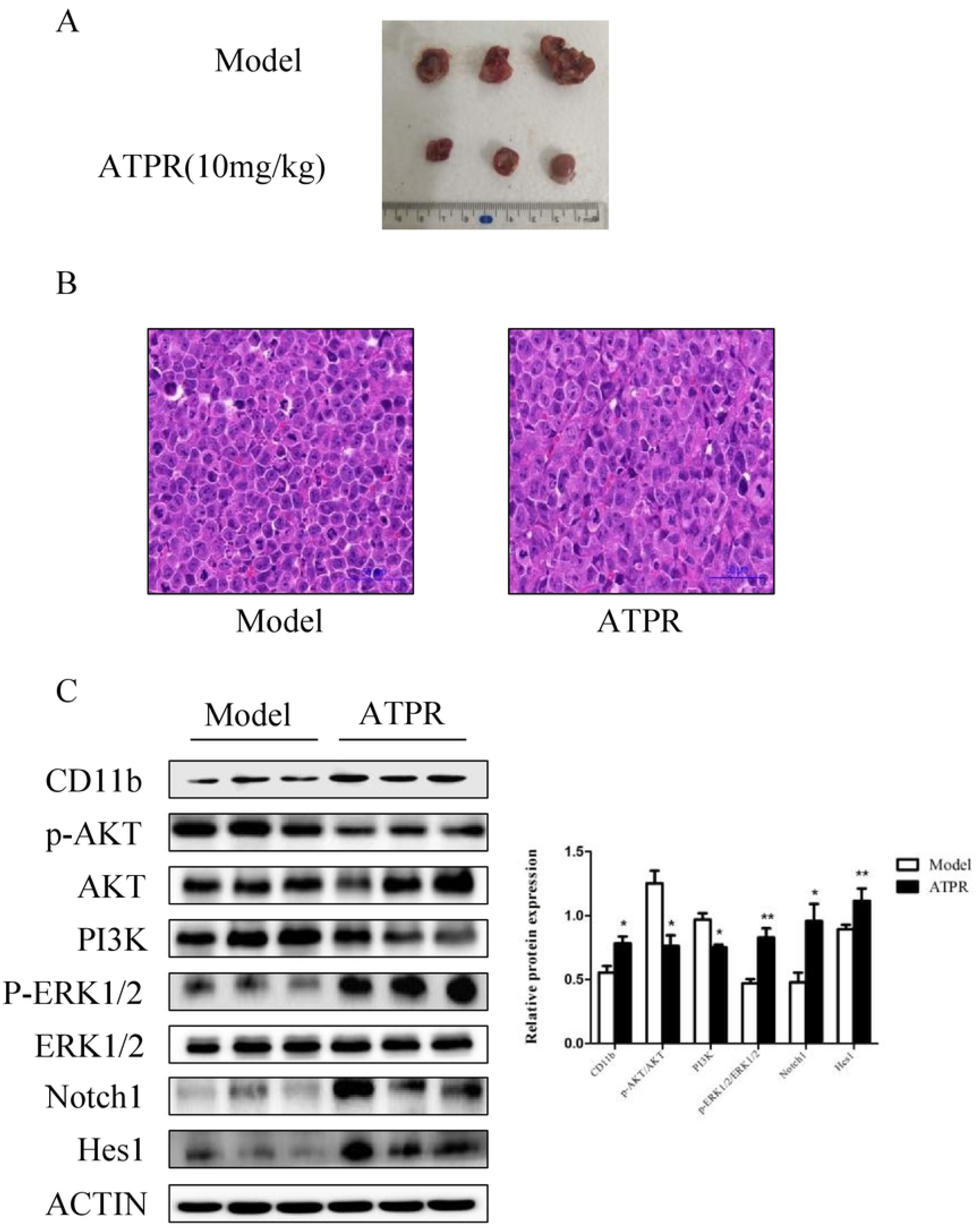
ATPR shows antileukemic effects *in vivo*. A, B. Pathology observation of Tumor tissues sections stained with HE (×400). C. Western blotting was performed to detect the expression levels of CD11b, p-AKT, AKT, PI3K, p-ERK1/2, ERK1/2, Notch1 and Hes1. \**p*<0.05, \*\**p*<0.01 versus control group.

## 4. Discussion

The NB4 cell line was derived from peripheral blood of patients with APL and undergoes differentiation induction by ATRA. Current clinical application of ATRA brings high proportion of CR for APL patients. But years of clinical observations showed that most of the initial treatment CR patients using ATRA alone occurred recent recurrence. Intensive chemotherapy was used to prevent relapse. Novel and effective inducers are wanted for further development of differentiation therapy [11]. ATPR, a derivative of ATRA, has been confirmed to repress proliferation and induce differentiation of various cancer cells effectively including breast, ovarian, esophageal, hepatocellular and gastric cancer [1]. Our previous studies have demonstrated that ATPR can inhibit proliferation, induce differentiation and apoptosis of NB4 [10], K562[17, 18] and SKM-1[19] *in vitro*. In the present study, we explored that the therapeutic effect of ATPR suppresses APL xenograft mouse model via the regulation of PI3K/AKT, ERK and Notch signaling pathway.

The PI3K/AKT, ERK and Notch signaling pathways play critical roles in the regulation of the cell apoptosis, proliferation, and differentiation in various cell types[20]. It has been reported that ATRA can induce vascular smooth muscle cells differentiation through rarα-mediated PI3K/Akt and ERK signaling pathways [16], and ATRA was found to play a role in restoring the insufficiency of mesenchymal stem cells via the Notch-1/Jagged-1 pathway[21]. Our results implicated that ATPR inhibits PI3K/AKT signaling activity but upregulates ERK and Notch signaling in APL-ascites mice models. The PI3K/AKT, ERK and Notch signaling pathway participated in ATPR treatment, and these pathways may play different roles in the process.

In conclusion, this study firstly demonstrates that ATPR can effectively inhibit tumor growth, induce cell apoptosis and differentiation via PI3K/AKT, ERK and Notch signaling pathways in the human xenograft mice model without obvious toxicity. These results will help us better understand the pathogenesis of APL and ATPR can potentially serve as a differentiation therapy against APL. Further studies may provide useful information for the development of a new therapeutic strategy against APL.

## Data availability statement

The data used to support the findings of this study are available from the corresponding author upon request.

## Acknowledgments

None.

